# NFC-enabled Sensing Platform for the Onsite Determination of Asparagine in Food

**DOI:** 10.1101/2025.07.14.664696

**Authors:** Hong Seok Lee, Ravleen Kaur Panesar, Laura Gonzalez-Macia, Giandrin Barandun, Firat Güder

## Abstract

Cooking foods rich in free asparagine (an amino acid that plays an important role in the synthesis of proteins), such as potatoes and flour-based products, can lead to the formation of acrylamide, a potent neurotoxin and potential carcinogen. Food manufacturers must quantify the levels of free asparagine before processing to minimize the formation of acrylamide, but conventional detection methods are often slow and expensive. We report a point-of-need measurement technology consisting of a chemically-functionalized paper-based electrical gas sensor and Near Field Communication (NFC)-enabled digital sensor interface (an integrated circuit) for rapid, cost-effective detection of free asparagine. Acid-functionalized paper-based electrical gas sensors selectively responds to ammonia gas, which is released during the enzymatic degradation of free asparagine. Through the use of the NFC-enabled digital sensor interface, a smartphone can wirelessly power and operate the paper-based gas sensor, enabling real-time quantification of free asparagine with a dedicated mobile application. We demonstrate a proof-of-concept sensor with a limit of detection of 3.19 µg/mL, capable of detecting low levels of free asparagine in food products. This practical tool can facilitate better control over the levels of acrylamide in food and improve food safety.

## Introduction

Staple foods such as potatoes, cereals, and legumes are essential components of the human diet, valued for their nutritional benefits and versatility. These staple crops naturally produce asparagine, a non-Sulphur containing amino acid that plays a central role in nitrogen metabolism. Asparagine functions as a safe, mobile form of nitrogen, facilitating both nitrogen storage and transport from source tissues to growing parts of the plant^1^. A portion of the asparagine in plant exists in its free (non-protein bound) form, known as free asparagine (FAsn). Plants often accumulate higher levels of FASn in response to environmental stresses such as drought, nutrient deficiency, or pathogen attack, and due to agronomic factors such as overfertilization with nitrogen, which increase nitrogen assimilation into free asparagine rather than protein. The accumulation of FAsn acts as a nitrogen reservoir to support growth and development under suboptimal conditions^1–3^. When foods rich in FAsn are cooked at a high temperature (above 120°C), the FAsn participates in the chemical reactions, known as the Maillard reaction, leading to the formation of acrylamide^4,5^.

Acrylamide is classified by the International Agency for Research on Cancer (IARC) as a probable human carcinogen, acrylamide poses considerable health risks, including neurotoxicity and carcinogenesis ^6–8^. National health authorities are establishing tolerable daily intake (TDI) levels for acrylamide: The European Food Safety Authority (EFSA) has indicated that dietary exposures exceeding 0.17 mg/kg of body weight per day may be associated with the growth of tumors and other adverse effects^9^. Especially in infants and children, acrylamide exposure poses a greater risk due to their lower body weight and higher dietary intake per kilogram of body weight compared with adults^10,11^. The European Commission has already set out legislation (EU 2017/2158) to control acrylamide levels in commonly consumed food products.^12^ The benchmark levels range from 40 µg/kg to 4000 µg/kg per kilogram of food product depending on the type of the product. Potato-based products (e.g., French fries, crisps) are regulated to have below 500-750 µg/kg, and cereal-based products (e.g., soft bread, breakfast cereals, biscuits) are regulated to have below 50-800 µg/kg of acrylamide ^8,13^. Food business operators are required to undertake representative sampling and analysis to comply with the food safety legislation aimed at reducing the presence of acrylamide in food products^14–16^. Addressing this potential health concern of acrylamide, it is recommended that acrylamide intake to be minimized as much as reasonably achievable.

To control acrylamide formation, efforts have primarily focused on reducing the concentration of its precursor, FAsn, in raw food products. Theoretical chemical mechanisms suggest that one mole of free asparagine can yield one mole of acrylamide via decarboxylation and deamination following Schiff base formation^17^. In processing conditions, however, the conversion efficiency is lower due to the complexity of food matrices and competing reactions^18^. Despite this fact, reducing the initial concentration of FAsn remains an effective strategy for lowering acrylamide formation during thermal processing as it is the molecular backbone for acrylamide. Strategies include breeding crop varieties with inherently lower asparagine content, enzymatic degradation of asparagine using asparaginase, and optimizing cooking parameters to limit generation of acrylamide.^19–21^. These strategies require optimizations to minimize formation of acrylamide while preserving the sensory and nutritional quality of the food, making systematic monitoring of FAsn levels essential for the food industry.

Current analytical methods for measuring asparagine, however, are not well suited for rapid, on-site industrial use. The gold standard, liquid chromatography-tandem mass spectrometry (LC-MS/MS), offers high sensitivity and specificity but requires expensive equipment and is confined to a laboratory ^22,23^. In contrast, colorimetric assay kits provide a faster and cost-effective alternative, but they still require a spectrophotometer and cannot be deployed outside of a lab^24^. Both methods suffer from being laboratory-bound, which leads to long turnaround times and high costs, hindering immediate decision-making during food processing^25^.

In this work, we report a point-of-need sensor system for on-site quantification of FAsn in food (**Figure 1**). For the measurement of levels of FAsn, we chemically functionalized paper-based electrical gas sensor (chemPEGS) and detected ammonia gas with high specificity, which is released during the enzymatic breakdown of asparagine in food samples. The sensor leverages Near Field Communication (NFC) technology, allowing for low-cost, user-friendly operation via a smartphone^26^. By simply positioning a smartphone above the sensor, the system wirelessly draws power and quantifies the concentration of asparagine in the sample through the dedicated mobile application.

**Figure 1.**
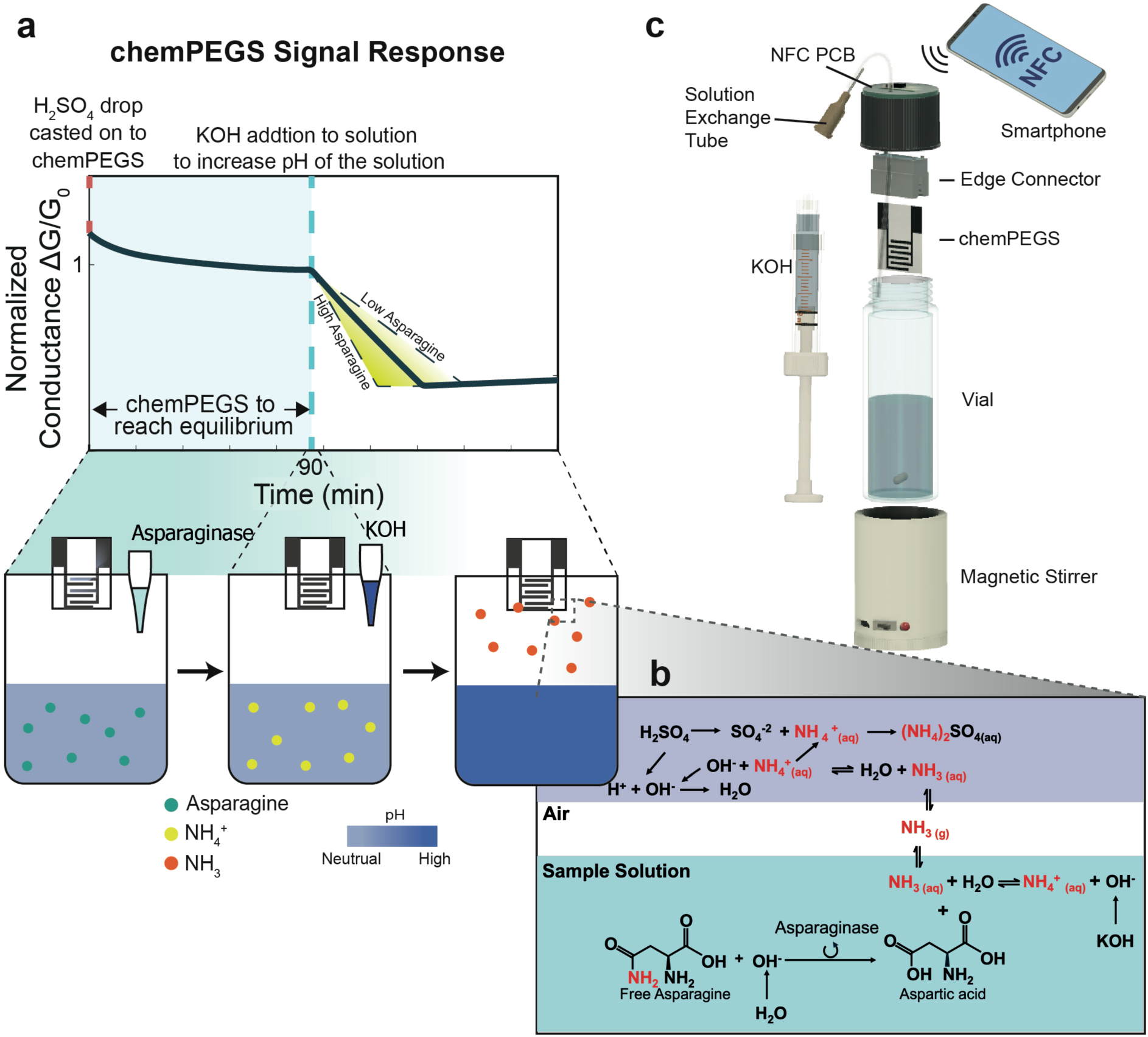
Scheme of the overall experimental procedure. **(a)** Changes in conductance of PEGS depending on the concentration of free asparagine in the solution when strong alkaline condition is introduced. **(b)** Free asparagine in the sample is broken down by asparaginase and produce NH_3_ (aq). Addition of KOH introduce strong alkaline conditions which causes NH_3_(aq) to volatilize to NH_3_(g) which subsequently arrives onto the fully humidified PEGS and neutralize the H_2_SO_4_. The neutralization reaction increases the ionic impedance thus decreases the conductance of PEGS. **(c)** The asparagine monitoring sensor kit consists of a 28.25ml glass vial with a modified lid to insert a disposable PEGS. On the top of the lid there is a NFC module to perform the measurement and communicate with a smartphone to operate the sensor.

## Results and Discussion

### Design and operation of NFC tag

We developed a compact NFC-based sensor tag designed to be mounted on the lid of the test vial for sensing NH_3_ in gas phase using chemPEGS. **Figure 2a** illustrates the block diagram of the electronic circuit, which consists of an energy harvesting module and the SIC4341 IC developed by Silicon Craft Technology PLC (Thailand). The NFC-based sensor tag harvests energy from the NFC antenna of a smartphone via inductive coupling with a planar copper coil antenna on the tag. The copper coil is tuned to resonate at 13.56 MHz, matching the internal capacitance of the SIC4341 (50 pF). The harvested energy is rectified by the internal rectifier of SIC4341 to generate a stable power supply. An external decoupling capacitor was added to the rectifier output to improve power stability. To ensure consistent operation, a low-dropout (LDO) regulator further stabilizes the harvested power, providing a constant 1.8V supply to the digital-to-analog converter (DAC), analog-to-digital converter (ADC), and potentiostat interface. An external 100 nF capacitor was connected to the output of the regulator to ensure reliable ADC performance.

**Figure 2.**
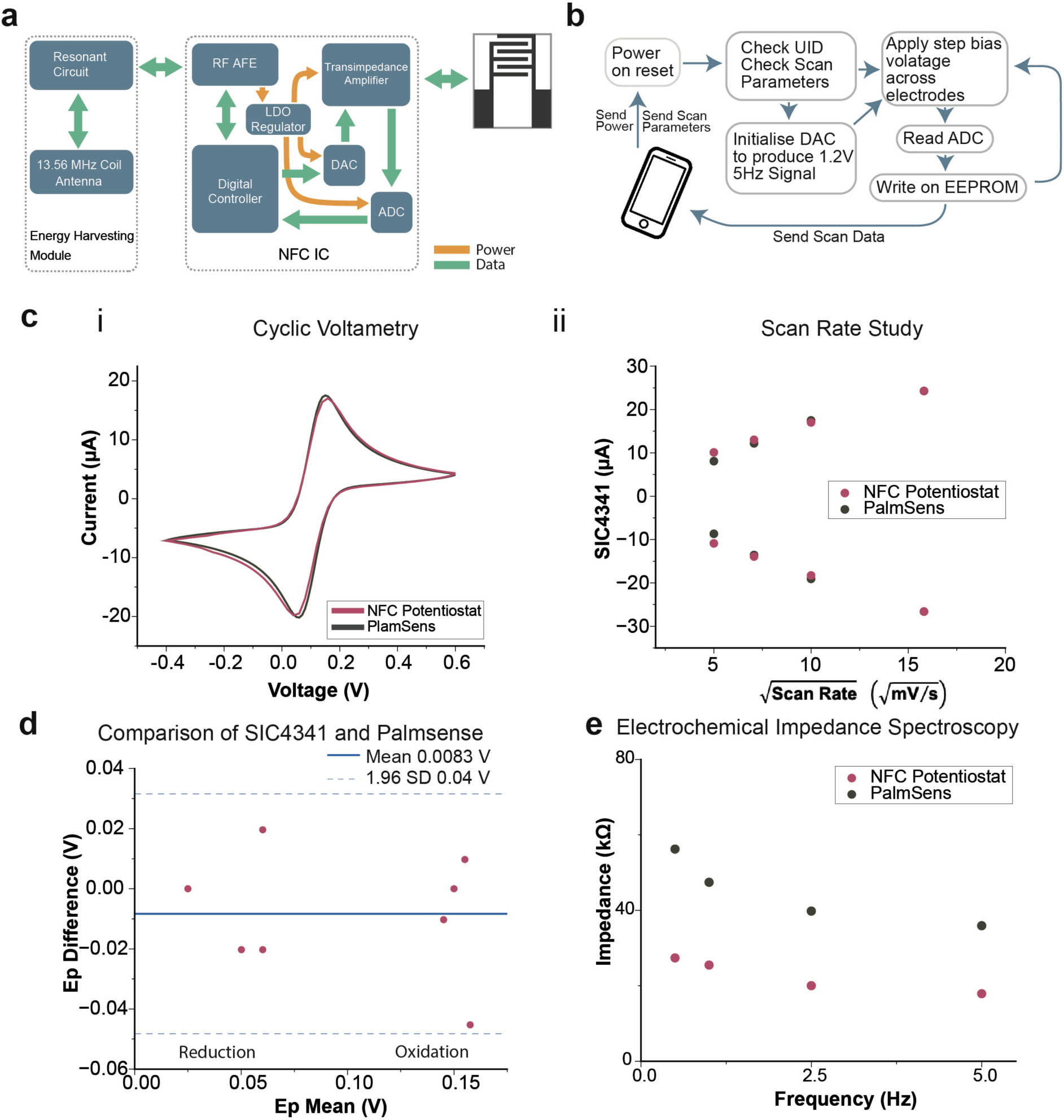
Operation and electrochemical characterization of the SIC4341-based potentiostat. a) Block diagram illustrating the key components of SIC4341 NFC circuit and connection to chemPEGS. b) Communication protocol between SIC4341 and a smartphone c) Comparison of the benchtop commercial potentiostat (PalmSens) to the NFC-based single chip potentiostat. i. Cyclic Voltammogram of 1 mM potassium ferricyanide at 50 mV/s; ii. Scan rate study: linear correlation of peak current and square root of scan rates as expected for diffusion-controlled processes d) Comparison of PalmSens and the single chip potentiostat by Bland-Altman plot of peak potentials (Ep) from the scan rate study. e) EIS comparison of the NFC and tabletop (PalmsSens) potentiostat

To communicate with the SIC4341 chip over NFC to perform measurements, we developed a custom-designed Android app using the Flutter open-source framework. Using the mobile app, the electrical signals applied to chemPEGS can be adjusted. The SIC4341 system allows the application of time-varying or fixed voltage signals to the paper-based sensor and measures the resulting electroanalytical currents. Once NFC communication is established between the smartphone and the tag, the chip is initialized, and the electrode potential is calibrated to nullify baseline noise between electrodes. This calibration step is crucial for optimizing the transimpedance amplifier and ADC settings, ensuring that only the current generated by the target electrochemical reactions is measured. Following calibration, the analyte solution is introduced, and the electroanalytical experiment is conducted. The current readings are digitized and saved in EEPROM of the chip. The reading is then forwarded from the sensor tag to the mobile phone over the NFC in real-time therefore the NFC data link must remain active throughout the measurement.

### Cyclic voltammetry (CV)

To validate the performance of the NFC-based single chip potentiostat (SIC4341), we compared its electroanalytical performance to a more advanced, benchtop commercial potentiostat (PalmSens4, PalmSens BV, The Netherlands) as shown in **Figure 2b**. The characterization was performed using commercially available screen-printed carbon electrodes (Metrohm DropSens) and 1 mM potassium ferricyanide in 0.1 M KCl. Before each measurement, fresh potassium ferricyanide solution was applied to all three electrodes. Between measurements, the electrodes were rinsed with deionized (DI) water and dried to ensure consistency.

Cyclic voltammograms were recorded at scan rates ranging from 25 to 2500 mV/s. The readings obtained using the SIC4341 were comparable to those measured with the commercial potentiostat, with both instruments producing similar peak shapes (Figure 2b and S1). Potassium ferricyanide concentrations exceeding 1 mM were not tested due to the ±20 μA current limit of the SIC4341. For the diffusion-controlled electrochemical reaction in 1 mM potassium ferricyanide, the peak currents were proportional to the square root of the scan rate, as expected for a diffusion-controlled electrochemical reaction. The peak currents and potentials recorded by both potentiostats showed strong agreement, with the NFC-based system exhibiting lower non-faradaic currents at scan rates below 50 mV/s. Bland-Altman analysis (**Figure 2c**) confirmed the agreement between the NFC potentiostat and the commercial system, with a mean difference of 8.34 μV. These results demonstrate that the NFC tag is fully capable of performing low-power, low-current electrochemical analysis, making it a viable alternative to conventional benchtop potentiostats for portable sensing applications in a disposable formfactor.

### Square wave voltage signal generation using the SIC4341

Our sensing system was based on the paper-based electrical gas sensor, which has previously been demonstrated to have high sensitivity for water-soluble gases when driven by a sinusoidal electrical signal^27^. To achieve a portable, battery-free system, we employed the SIC4341 IC to generate a square wave signal across the two electrodes through a set of customizations to the software (Figure S2). A square wave was selected due to the limitations of the IC chip, which imposes a minimum conversion time of 100 ms to digitize the analog signals. This limitation exists because the IC can only assign a new accurate bias voltage every 100 ms. The half-duplex nature of the NFC protocol (ISO 14443A) creates this inherent communication delay^28^. The protocol mandates sequential communication, where one device must wait for the other to complete its transmission before initiating a response. This constraint also makes generating a continuous, high-frequency sinusoidal signal unattainable. To address this constraint, SIC4341 was programmed to output a square wave, effectively compensating for the extended conversion time. The maximum achievable square wave frequency with the NFC tag developed was 5 Hz, with the working and counter electrode voltages alternating every 100 ms.

Upon receiving power and scan parameters from a smartphone, the SIC4341 Power-On-Reset (POR) is triggered, resetting all internal logic and registers to their default state before storing the user-defined parameters in the EEPROM non-volatile memory (Figure 2b). These parameters include electrode pin configuration, voltage bias values, duration for each biasing step, and ADC sampling rate. An on-chip 8-bit DAC sets the voltage levels for each electrode, with an assignable range between 0.4 V and 1.6 V, generating a 1.2 Vpp square wave signal (−0.6 V to +0.6 V). This is achieved by alternating the voltages of the working electrode (V_WE_ = 1.2 V) and the reference electrode (V_RE_ = 0.4 V). After 100 ms, V_WE_ and V_RE_ switch values, producing a square wave, as shown in Figure 2c. The current flow through the working electrode is measured using the built-in transimpedance amplifier of the chip, with signal conversion performed via ADC sampling during each voltage cycle.

Electrochemical impedance spectroscopy (EIS) confirmed that the 5 Hz square wave signal is suitable for electrochemical measurements (**Figure 2e**). At 100% relative humidity (RH), the impedance of chemPEGS plateaus, indicating minimal electrode-electrolyte capacitive behavior and revealing the intrinsic resistance of the fully humidified paper substrate. Any changes in the conductivity of the substrate (paper) can therefore be attributed to the presence of the target analyte, validating the effectiveness of the NFC-powered electrochemical sensor.

### Quantifying Free Asparagine using chemPEGS

Asparagine is a water-soluble amino acid that can be simply extracted from food samples by dispersing them in distilled water. FAsn extracted can be detected by exploiting an enzymatic reaction. In the presence of asparaginase, FAsn breaks down into aspartate and produces ammonia (equation 1).

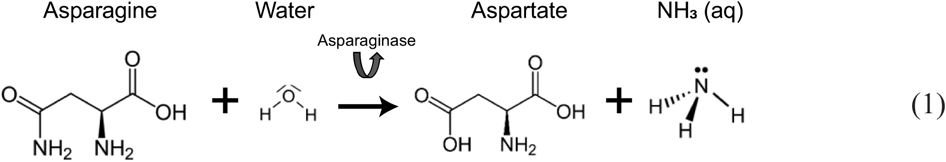

The ammonia in solution exists in equilibrium with ammonium ions, as shown in equation 2. This equilibrium is largely influenced by temperature and the solution pH.

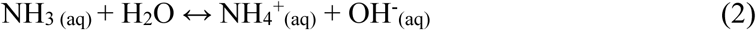

At room temperature, increasing the pH of the solution by the addition of a strong base shifts the equilibrium to the left, converting NH_4_^+^_(aq)_ to NH_3_(aq) according to Le Chatelier’s Principle. The relationship between pH and the ratio of NH_3 (aq)_ to NH_4_^+^_(aq)_can be further explained using the Henderson-Hasselbalch equation^29^:

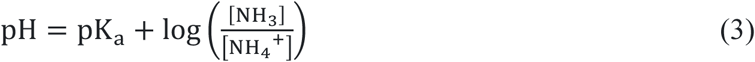

When the pH exceeds the *pK_α_* of ammonium (approximately 9.25 at 20°C), NH_3 (aq)_ becomes the dominant species. At pH > 11, nearly all NH_4_^+^_(aq)_ions are converted to NH_3(aq)_, which volatilize into the gaseous form (NH_3 (g)_), leaving the sample solution and entering the headspace of the testing vial^30^.

Paper, made of highly hygroscopic cellulose fibers, adsorbs a substantial amount of moisture from the surroundings in a closed humidified system and form a thin layer of water on the surface of the paper^31,32^. In the presence of volatile water-soluble gases, these gases can interact with the water adsorbed in paper. If the gaseous species in the closed system are water soluble (such as ammonia), they dissolve in the layer of water and dissociate to become ions which consequently alter the electrical impedance of paper due to a change in ionic conductivity^33^. By printing two electrodes on paper, the change in electrical impedance due to water-soluble gases can be detected^34^. To make the paper-based sensor more specific to NH_3_, H_2_SO_4_ was deposited onto the sensing area – between two electrodes by adding 5μl of 0.03M. The acid introduces two additional hydrogen ions [H^+^] and a sulphate ion [SO_4_^-2^]) to the water layer formed on the paper substrate. As the NH_3_ arrives in the layer of water, NH_3_ dissociates into ammonium and hydroxide ions (NH_3_ + H_2_O → NH_4_^+^ + OH^-^). The OH^-^ ions in the solution layer are neutralized by the existing H^+^ ions and generate neutral water molecules (H^+^ + OH^-^ → H_2_O). Subsequently, the remaining NH_4_^+^ and SO_4_^2-^ ions become highly soluble salts (ammonium sulphate) which once again dissolve inside the water layer. The resulting chemical reaction substitutes the H^+^ ion with NH_4_^+^. As the mobility of the ammonium ion in solution is lower than the mobility of the hydrogen ion, the overall electrical impedance of the paper increases (**Figure 1**)

We first validated our sensor system with our custom built transimpedance amplifier circuit with a gain matching the transimpedance amplifier of SIC4341 (Figure S3). The electrical measurements were collected through an Arduino Due board and processed in MATLAB (Figure S4). 1.6 V_pp_ signal was applied for the characterization against different NH_4_OH concentrations (62.5, 312.5, 625, 3125, 6250 mM). All experiments were performed with new chemPEGS, and the test solutions were stirred throughout. chemPEGS was inserted into the custom designed lid with an electrical feedthrough (**Figure 3d**). 16 mL of test solution was added to the standard 28.25 mL glass vial (VWR) for each measurement, as this volume ensures that the chemPEGS remains unaffected by direct contact with the test liquid, even if the vial is subjected to gentle knocks during handling. This is crucial to maintain the integrity and functionality of the sensor, as contact with the solution would compromise its performance. Additionally, 16 mL of test solution minimizes the headspace, leaving only 12.25 mL within the vial, which is critical as it reduces the dilution of ammonia gas in the headspace. A smaller headspace leads to a higher concentration of ammonia gas, which enhances the sensitivity of the chemPEGS to detect the gas effectively. This careful consideration of the volume of the test solution was therefore essential for achieving reliable and reproducible measurements. The solution exchange tube is also submerged beneath the test solution so that no air is leaking out through the tube to maintain the closed system. The tube is later used to extract a portion of test solution using a syringe, and the same volume is immediately replaced by injecting 2M KOH to increase the pH of test solution without disturbing the equilibrium of the liquid-vapor inside closed vial.

**Figure 3.**
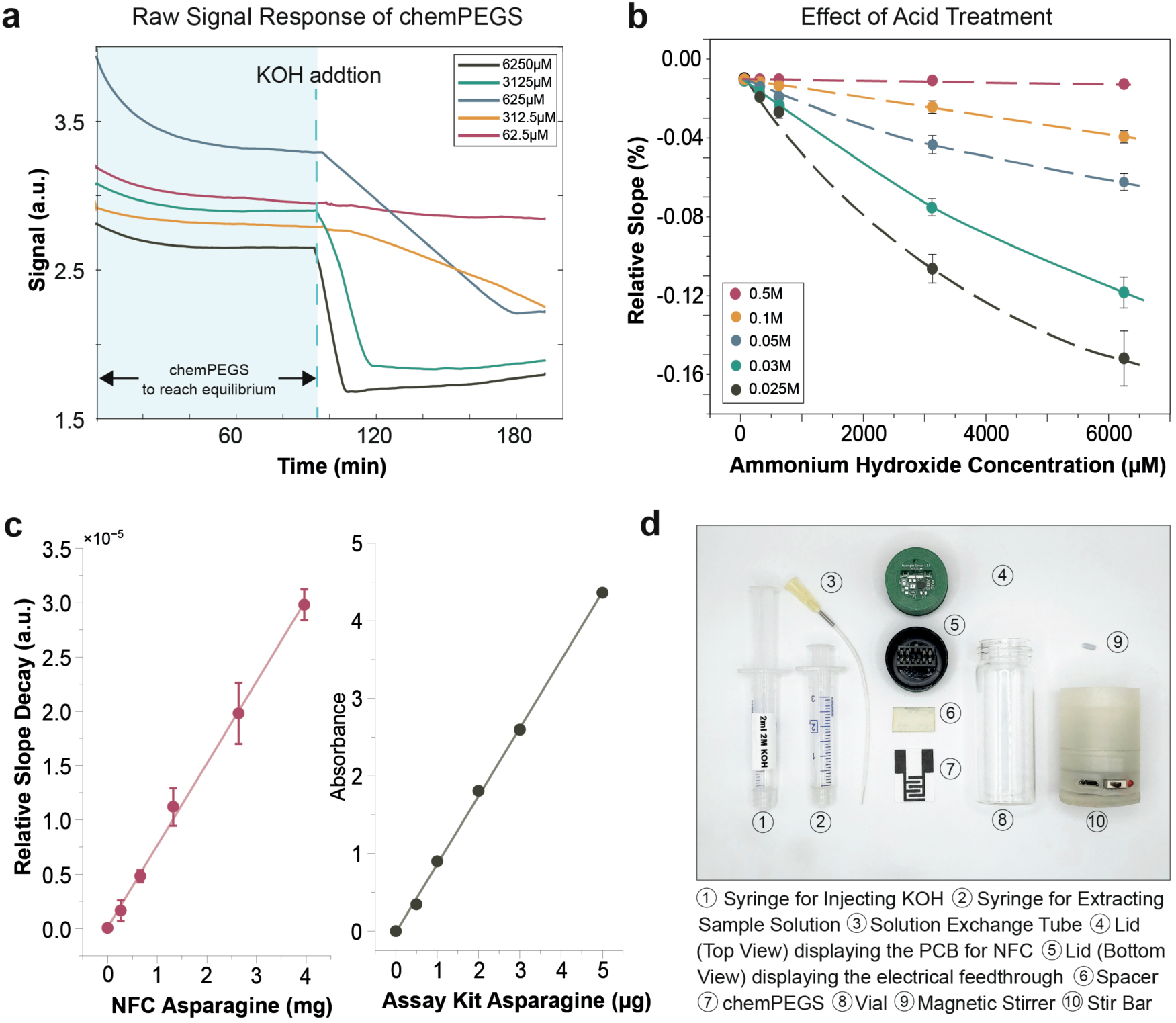
**(a)** Raw signals acquired from the chemPEGS at different concentrations of NH_4_OH (62.5, 312.5, 625, 4125, 6250 µM. Higher concentration of NH4OH causes a steep drop in signal **(b)** The relative drop in signal, ΔG/G_0_ was plotted against the H_2_SO_4_ concentrations (0.5, 0.1, 0.05, 0.025 M) drop casted on the paper-based electrical sensor (n = 5). Lower concentration of H_2_SO_4_ exhibits a larger drop in signal as acid is neutralized more quickly with NH_3_. **(c)** Calibration curve of the developed NFC-based FAsn sensor kit and the commercial colorimetric assay kit (n = 5). **(d)** Photograph of the NFC-based FAsn sensor

When the chemPEGS is fully humidified and its conductance is stabilized, 2ml of test solution was replaced with 2M KOH. The pH of the test solution increased to 14 and the ammonia equilibrium shifted toward NH_3_ and suppressing NH_4_^+^. The NH_3_ volatilized into the headspace and reached the sensing area of the chemPEGS, resulting in a steady decline in conductance (**Figure 3a**). This decline in conductance continued until all the deposited H_2_SO_4_ was neutralized by ammonia. Once the acid was fully neutralized, the conductance began to increase due to the presence of excess ammonia, which contributed to a rise in ionic conductivity. It was observed that the rate of decay for the signal (conductance) after the addition of KOH was directly proportional to the concentration of ammonium ions in the test solution. To account for variations arising from inconsistencies associated with fabrication and differences in material properties among individual chemPEGS, the conductance of each sensor was normalized against its G_0_ (chemPEGS conductance immediately prior to the addition of KOH to the test solution). This normalization ensured that observed trends reflected the true sensing behavior of the system, not artifacts from manufacturing discrepancies or inhomogeneity of the paper substrate.

The calibration curves shown in Figure 3b include measurements conducted at varying concentrations of H_2_SO_4_ applied to the chemPEGS. Increasing the concentration of H_2_SO_4_ broadens the dynamic range for sensing while simultaneously reducing the sensitivity of the sensor. This behavior is attributed to the shift in the ratio of H^+^ to neutral molecules at a fixed concentrations of ammonia in the headspace of the vial. At higher concentrations of H_2_SO_4_, a larger quantity of acid ions is deposited on the chemPEGS, leading to a scenario where only a small fraction of these ions is neutralized upon exposure to ammonia. As a result, the un-neutralized H^+^ dominates the overall ionic conductivity, which diminishes the relative sensitivity of the chemPEGS due to a slower decline in conductivity. Conversely, at lower concentrations of H_2_SO_4_, the ratio of ions to neutral molecules shifts more significantly upon ammonia exposure. Each neutralization induces more pronounced reduction in ionic conductivity, resulting in steeper changes in the sensor’s response. Among the tested concentrations, 0.03 M H_2_SO_4_ demonstrated the optimal balance between dynamic range and sensitivity for measuring free asparagine in solutions originating from food products. chemPEGS functionalized with 0.03 M H_2_SO_4_ is, therefore, the most suitable condition for detecting ammonia generated from the enzymatic degradation of asparagine.

### Measuring concentrations of FAsn in food samples

The NFC tag developed in FAsn sensor kit was used for testing with real food samples. The sensor kit was calibrated to exhibit linear performance for concentrations for FAsn up to 30 µmol per 16mL, with a limit of detection (LOD) of 3.19 µg/mL as shown in **Figure 3c**. The detection performance of the sensor was cross-validated against a commercially available L-asparagine assay kit (Megazyme). A variety of potato samples (Red, White, and Maris Piper) were tested. The samples were centrifuged to remove dense particles such as starch. For the commercial assay kit, the samples were further deproteinized to avoid interference with spectrophotometric measurements.

The Bland-Altman plot confirmed agreement between the two methods, as all values fell within the limits of agreement (±1.96 SD), with a consistent positive bias in chemPEGS sensor relative to the commercial test (mean = 158.9 µg of asparagine per gram of potato). This bias originates from the hydrolysis of proteins in potatoes under strong alkaline conditions, which could generate additional ammonia as a background signal and affect the conductance of the chemPEGS. The gradual breakdown of proteins into smaller peptides and amino acids during hydrolysis likely led to the release of ammonia over time, contributing to change in signal of the chemPEGS.

In samples without asparaginase, however, no significant drop in signal was observed, particularly in the early stages of the experiment (**Figure 4**). The effect of protein hydrolysis on the conductance of chemPEGS appeared more prominent after 20 minutes following the addition of KOH. As the rate of change in conductance was measured within the first 5 minutes of the addition of KOH, errors arising from the hydrolysis of proteins were minimized.

**Figure 4.**
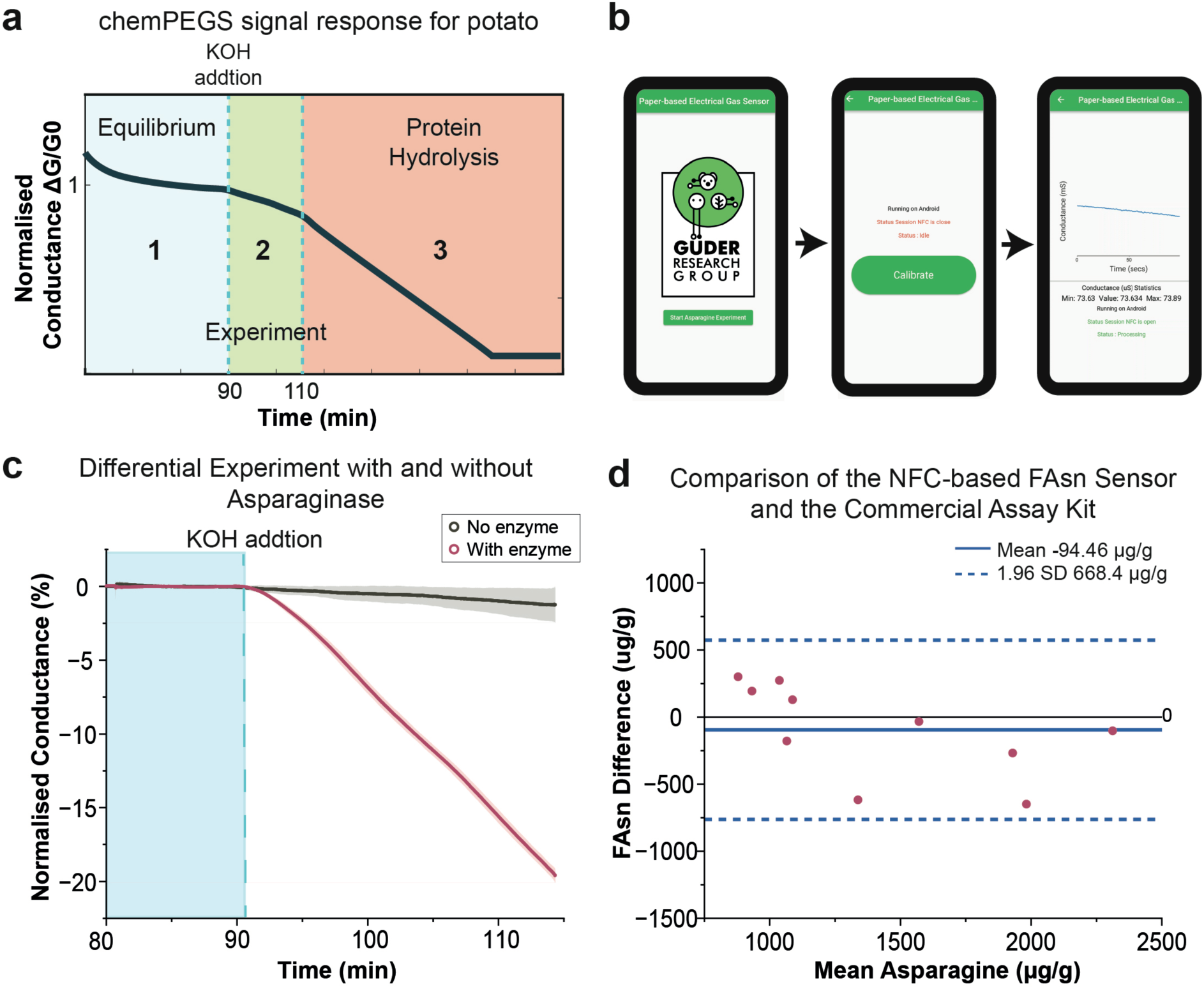
**(a)** Illustration of change in relative conductance of chemPEGS throughout the experiment: 1. Conductance reaches equilibrium at around 90 min 2. KOH addition causes ammonia from test solution to reach the chemPEGS and conductance drops 3. Further drop occurs after 20min for the KOH addition due to proteins hydrolysis which also generate ammonia gas. **(b)** Screenshots of the developed Android mobile app consists of three main stages 1. Powering stage 2. Calibration 3. Data Collection. **(c)** Differential experiment with and without asparaginase enzyme (n=3). **(d)** Bland-Altman plot comparing measurements of FAsn in potato samples between the developed NFC-based FAsn sensor and commercial FAsn assay kit by Megazyme. The solid blue line represents the mean difference (bias) of −94.46 µg/g. The dashed blue lines represent 95% limits of agreement (n = 10)

## Conclusion

The FAsn sensor kit developed in this work requires minimal laboratory equipment, needing only a centrifuge for sample preparation, unlike conventional methods that rely on specialized instruments such as high-performance liquid chromatography (HPLC) systems and spectrophotometers. The entire sensing process, including signal generation, measurement of electrical signals, and data transmission, is managed by the SIC4341, which operates wirelessly through a smartphone without the need for additional batteries. The developed sensor kit demonstrated high accuracy for FAsn detection (R² = 0.99) and is suitable for measuring asparagine levels in food samples like potatoes, following appropriate sample preparation. This system enables on-site monitoring of FAsn levels in food processing facilities and farms, significantly reducing testing costs (US $1.39 per test) (**Figure. S5**) and analysis turnaround time. This allows raw ingredient suppliers to take timely and appropriate actions to reduce FAsn levels before cooking, thereby mitigating high levels of acrylamide in food products during processing.

Beyond industrial and research applications, this sensor could also be adapted for home use by health-conscious consumers or individuals requiring a strict asparagine- controlled diet^35^. By providing a low-cost, portable, and battery-free solution for the detection of free asparagine, this sensor kit has the potential to enhance food safety, reduce acrylamide exposure, and improve quality control processes across various sectors.

Despite its advantages, the sensor kit developed presents three key challenges: (i) The wireless communication between the smartphone and NFC sensor tag must remain stable throughout the measurement process. Movements can interrupt data transmission and delivery of power, leading to incomplete results. Integrating a secure phone mount into the sensor kit would help ensure consistent and reliable NFC communication. (ii) chemPEGS requires encapsulation with a membrane to prevent direct contact with the test solution, as physical disturbances can cause splashes, potentially damaging the sensor. A viable solution would be to incorporate gas-permeable membranes, such as those reported by Naik et. al^36^. A gas permeable membrane would only allow gases and vapors to pass through to the sensor while blocking contact with liquids, eliminating the possibility of contaminating of the sensor and increasing stability^37^. (iii) The strong alkaline conditions used during the measurements may cause denaturation and hydrolysis of proteins, affecting the integrity of the food sample. Other food components, such as carbohydrates, lipids, or vitamins, may degrade and produce by-products that could interfere with the readings. Although this was not an issue in our experiments involving potatoes, liquid samples with high amounts of reducing sugars or water-soluble vitamins (vitamin B, C) might exhibit more pronounced effects. Implementing differential measurements, where readings are taken with and without the asparaginase enzyme, could help eliminate unwanted background responses.

While this study primarily focuses on measuring FAsn in potatoes, the system is highly adaptable and could be used for various food samples requiring strict monitoring of FAsn with appropriate modifications to sample preparation. The sensing system can also be further optimized to reduce test time by adjusting parameters such as the size of chemPEGS and volume/concentration of H₂SO₄ applied, allowing chemPEGS to reach equilibrium faster. Additionally, the NFC-enabled paper-based sensing platform could be expanded for detecting other analytes, such as urea in blood, by measuring ammonia released from enzymatic urease reactions^38^. The technology reported in this work demonstrates the feasibility of portable, low-cost, and NFC-powered electrochemical sensing that can be used at the point-of-need, paving the way for its integration into monitoring food safety, diagnostics for human and animal diseases, and real-time environmental analysis.

## Methods

### Asparagine sensor kit fabrication

All experiments were conducted using a standard 28.25 mL glass vial (VWR). A 12-position female dual-edge connector (Part number: 7-5530843-5, TE Connectivity) was modified to function as a feedthrough for integrating the chemPEGS with external electronics. The connector’s contact pins were soldered to wires, allowing connection either to a transimpedance amplifier circuit configured with a 128 kΩ gain resistor for chemPEGS characterization or directly to the sensor I/O pins of the NFC chip in the final integrated design. The NFC tag PCB (28 mm diameter) was designed using KiCAD and manufactured by JiaLiChuang Co., Ltd. The SIC4341 chip (Silicon Craft Technology PLC) provided a potentiostat sensor interface along with NFC capabilities, enabling both energy harvesting and wireless communication with a smartphone. Passive components, including two 100 nF capacitors, were sourced from Digi-Key Electronics. For controlled chemical solution exchange, a silicone capillary tube (1 mm inner diameter, 1.5 mm outer diameter) from RS Components was incorporated into the vial cap. A blunt 18G Luer lock syringe needle (VWR) was attached to the end of the capillary tube to ensure secure and precise solution transfer. The capillary tube and connector were sealed to the vial cap using a glue gun to maintain an airtight system. Solution exchange was performed using a 2 mL Luer lock syringe (VWR).

### Chemically Functionalized Paper-based electrical gas sensor (chemPEGS) fabrication

The chemPEGS were fabricated using a screen-printing method on Whatman Chromatography 1 cellulose paper (General Electric Healthcare). To enhance printability, carbon ink (C2030619P4, SunChemical) was diluted with a proprietary diluent (S60118D3, Gwent Electronics) at a 70:30 (w/w) ink-to-solvent ratio. After printing, the sensors were cured in an oven at 40 °C for 60 minutes to remove excess organic solvents from the electrodes. The average resistance of each electrode was measured to be 5.34 ± 0.72 kΩ. Prior to each experiment, 5 μL of 0.03 M sulfuric acid (Sigma Aldrich) was drop-cast onto the paper to optimize sensor performance.

### Characterization of the Asparagine Sensor Kit

A 5 Hz, 1.6 V peak-to-peak signal was applied to one of the chemPEGS electrode, and its impedance response was continuously measured using the developed asparagine sensor kit. The kit was connected to a custom-built transimpedance amplifier circuit, which interfaced with an Arduino Due for signal processing. Initial calibration of the asparagine sensor was performed using NH₄OH solutions (Sigma Aldrich) with concentrations of 62.5, 312.5, 625, 3125, and 6125 μM. Additionally, different concentrations of H₂SO₄ (Sigma Aldrich)—0.5, 0.1, 0.05, and 0.025 M—were deposited onto the chemPEGS to investigate the sensor’s response. Each vial was filled with 16 mL of DI water, and 5 μL of H₂SO₄ was drop-cast onto the sensing area of chemPEGS just before sealing the glass vial. The addition of acid facilitated the achievement of 100% RH equilibrium more rapidly due to its evaporation into the surrounding environment. After 1.5 hours, the system reached conductivity equilibrium (G₀), where no further change in the signal was observed over time (Figure 3a). To introduce the test solution, a portion of the water was removed via the solution exchange tube, and concentrated NH₄OH was added to obtain the desired concentration. This step was necessary to prevent baseline inaccuracies, as highly concentrated NH₄OH solutions are inherently basic, causing some NH₄⁺ to leave the solution. Subsequently, 2 mL of the test solution was withdrawn, and 2 mL of KOH was introduced using a syringe. This process increased the pH of the test solution while maintaining equilibrium conditions.

### FAsn Extraction from potato

Potato skins were peeled, rinsed with water, and ground into small pieces. 20 g of the processed potato was dispersed in 100 mL of deionized (DI) water and stirred for 30 minutes. The mixture was then strained using a fine strainer, and additional juice was extracted by pressing the pulp with the back of a spoon. The collected potato juice was centrifuged at 1500 × g for 10 minutes, and the resulting supernatant was used for the experiment.

### L-asparagine assay kit

The L-Asparagine/L-Glutamine/Ammonia (Rapid) Assay Kit by Megazyme was used to cross-validate the performance of developed sensor. The method involves enzymatic reactions followed by spectrophotometric detection. 15 mL of extracted potato juice was mixed with 5 mL of ice-cold 1 M perchloric acid (Sigma Aldrich) and centrifuged at 1500 × g for 10 minutes. The resulting supernatant was neutralized with 2 M KOH (Sigma Aldrich) to adjust the pH to 7–9, the optimal range for asparaginase activity. pH adjustments were monitored using a Hanna Instruments H15222 benchtop EC/pH meter. 15 mL of extracted potato juice was mixed with 5 mL of ice-cold 1 M perchloric acid (Sigma Aldrich) and centrifuged at 1500 × g for 10 minutes. The resulting supernatant was neutralized with 2 M KOH (Sigma Aldrich) to adjust the pH to 7–9, the optimal range for asparaginase activity. pH adjustments were monitored using a Hanna Instruments H15222 benchtop EC/pH meter. 10 mL of the neutralized supernatant was placed in 96-well microplate (Corning Inc.) and combined with 20 mL of buffer (pH 4.9) and 2 mL of glutaminase. The mixture was incubated for 5 minutes at room temperature to convert L-glutamine to glutamate and ammonia. Afterward, buffer (pH 8.0), NADPH, and water were added, and incubated for another 5 minutes. Glutamate dehydrogenase (2 μL) was then added to catalyze the reaction between ammonia and 2-oxoglutarate, and the first absorbance was measured using the microplate reader (Thermo Scientific). Finally, asparaginase (2 μL) was added to hydrolyze L-asparagine to L-aspartate and ammonia, and the second absorbance was taken after 5 minutes. The decrease in absorbance correlates with NADPH consumption, enabling quantification of FAsn using the extinction coefficient-based formula.

### Characterization of the Developed FAsn Sensor Kit

FAsn solutions with concentrations of 0, 125, 312.5, 625, 1250, 1875 μM were prepared by dissolving L-asparagine (Sigma Aldrich) in deionized (DI) water. A separate asparaginase solution was prepared by dissolving asparaginase from *Escherichia coli* (Sigma Aldrich) in distilled water to achieve a final concentration of 16 units/mL. (One unit of asparaginase catalyzes the release of 1 μmol of NH₃ from L-asparagine per minute at pH 8.6 and 37 °C.)

For each experiment, 16 mL of the FAsn solution and 50 µL (0.8 units) of asparaginase were added to a vial. The chemPEGS was placed inside the feedthrough, and the vial was sealed. The vial was positioned on a custom-designed magnetic stirrer set to 400 rpm, and the solution was continuously stirred throughout the experiment. The magnetic stirrer was specifically designed to generate a low magnetic field to prevent interference with NFC communication. The system was left for 90 minutes to allow the chemPEGS to reach equilibrium and for asparaginase to fully degrade the FAsn in the test solution.

Next, the Android app was launched on a smartphone, which was positioned above the sensor. The smartphone began recording the signal (current) as 2 mL of the test solution was withdrawn via the solution exchange tube and replaced with 2 mL of 2M KOH. The collected data was subsequently transferred to a computer for analysis.

### App Design for a Smartphone

An Android app was developed by modifying the SIC4341 library from SiliconCraft Ltd. using Flutter. The app controls the generation of specific bias voltages on each electrode over a predefined time period and measures the current on the working electrode. The app operates in three main stages: 1. Powering stage: The NFC tag receives power and initiates communication between the smartphone and the sensor. 2. Calibration: The potentiostat interface and ADC is calibrated to ensure accurate measurements 3. Data Collection: The user can monitor conductance in real-time on the smartphone screen (**Figure 4**).

## Acknowledgments

The authors thank the Department of Bioengineering at Imperial College London and SiliconCraft Ltd for providing SIC4341 IC chips and their application library. F.G. and H.S.L. thank the Economic and Social Research Council (ES/P000703/1), and London Interdisciplinary Social Science Doctoral Training Partnership. L.G.-M. thanks the European Union’s Horizon 2020 research and innovation program under the Marie Sklodowska-Curie grant agreement no. 101025390. F.G. and G.B. thank the Engineering and Physical Sciences Research Council (EPSRC, EP/R010242/1), and General Electric Healthcare. F.G. thanks Bezos Earth Fund through the Bezos Centre for Sustainable Protein (BCSP/IC/001).

