## Supplementary Information for "NFC-enabled Sensing Platform for the Onsite Determination of Asparagine in Food"

**NFC-enabled Paper-based Sensor for the Detection of Asparagine in Food**

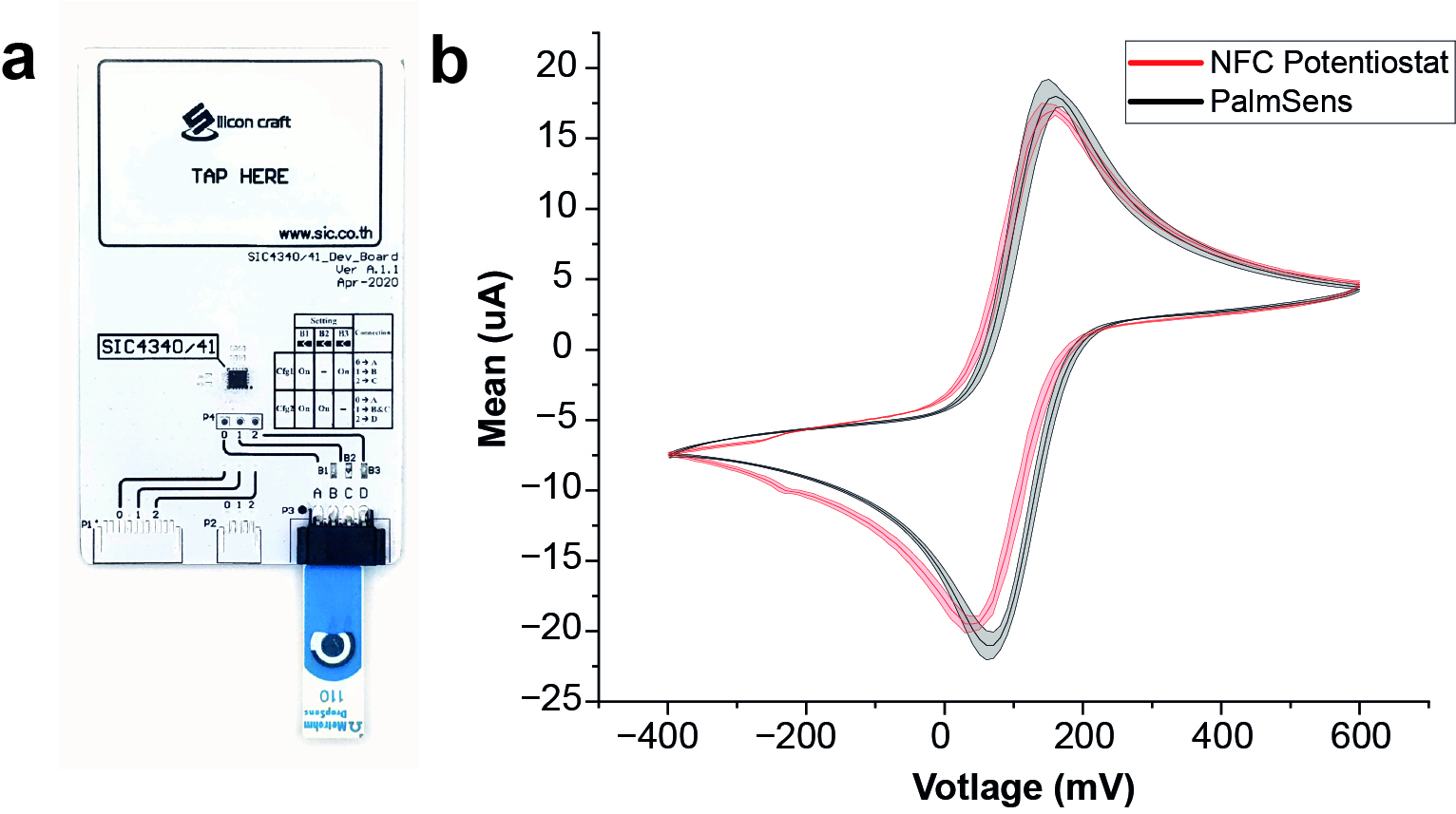

**Figure S1**. **(a)** Photograph of the single chip NFC-based potentiostat **(b)** Cyclic voltammogram comparison of the benchtop commercial potentiostat (PalmSens) to the NFC-based single chip potentiostat using1mM Potassium Ferricyanide in 0.1M KCl at 100mV/s (n = 5)

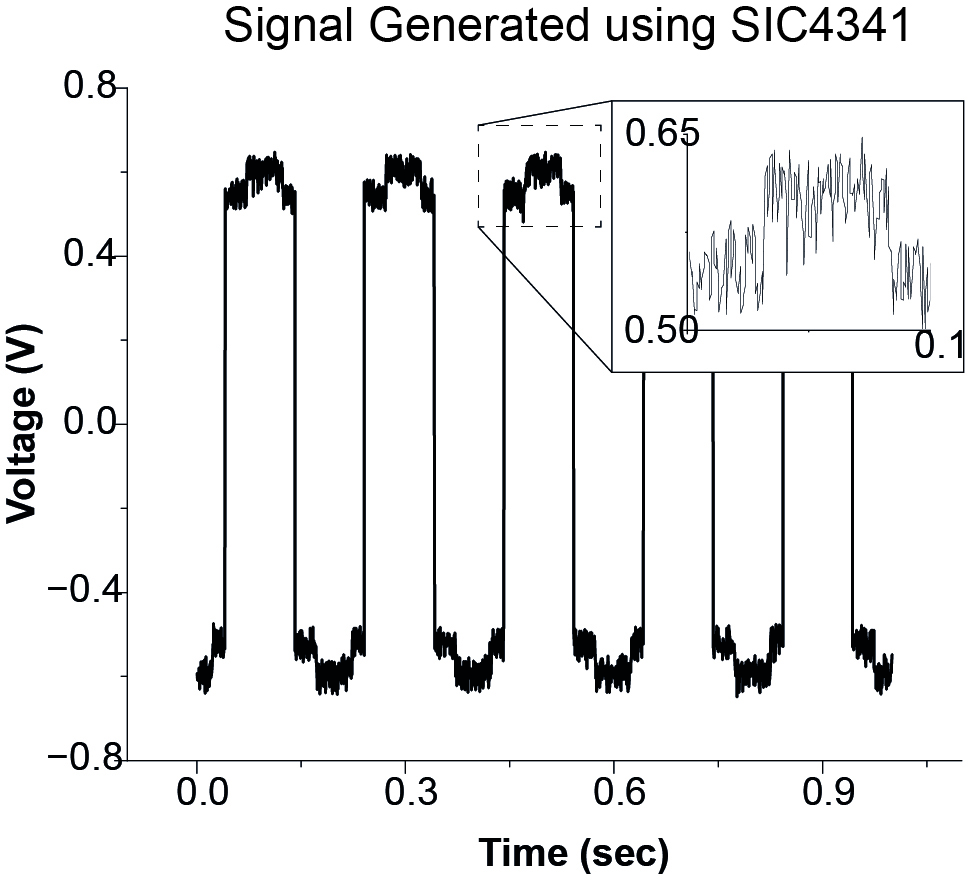

**Figure S2**. Generation of a 5Hz, 1.2V peak-to-peak square wave using the SIC4341 IC, which was commanded by the developed smartphone app. The chip applied an alternating potential (swinging between +0.6V and -0.6V) to the electrodes every 100ms. The resulting signal was applied to the chemPEGS.

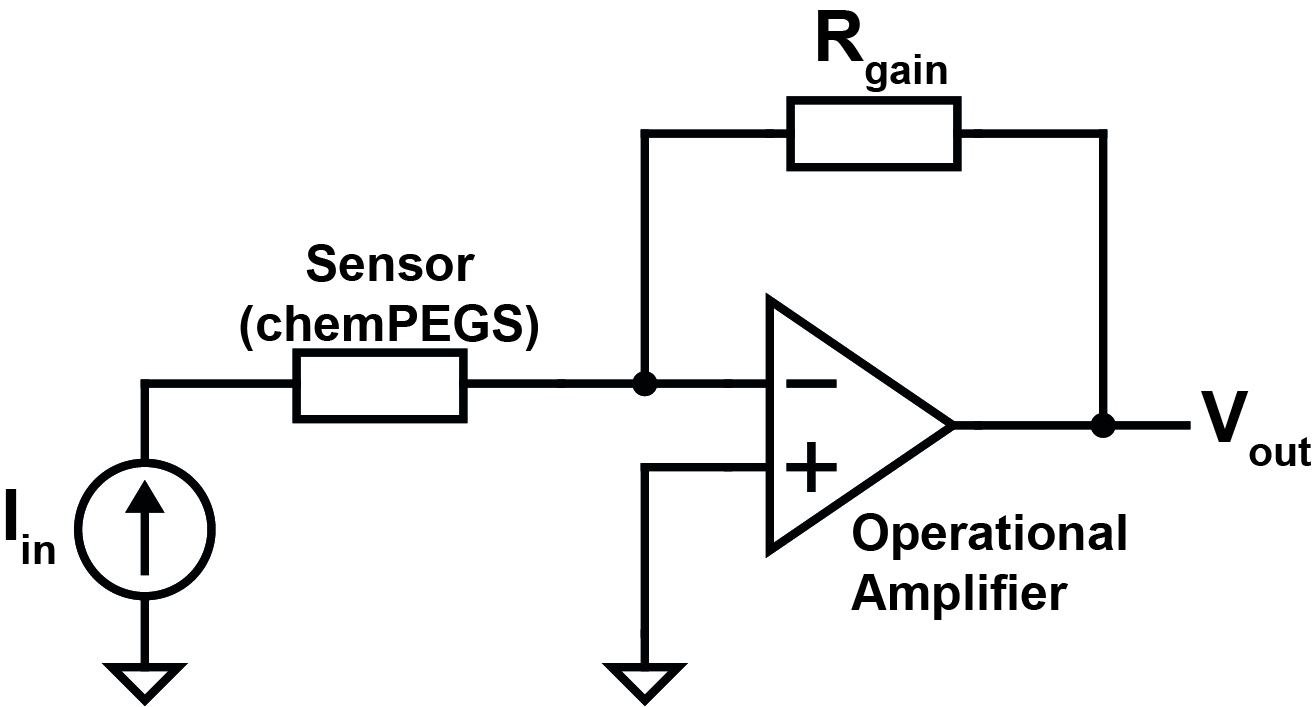

**Figure S3**. To measure the ionic impedance of the chemPEGS, a transimpedance amplifier converts the sensor's current (I_in_) into a proportional voltage (V_out_). The amplification factor is set by the gain resistor (R_gain_), following the relationship:

V_out_ = – R_gain_ × I_in_

The resulting analog voltage (V_out_) is read and digitized by the built-in ADC of the SIC4341 IC or the Arduino Due.

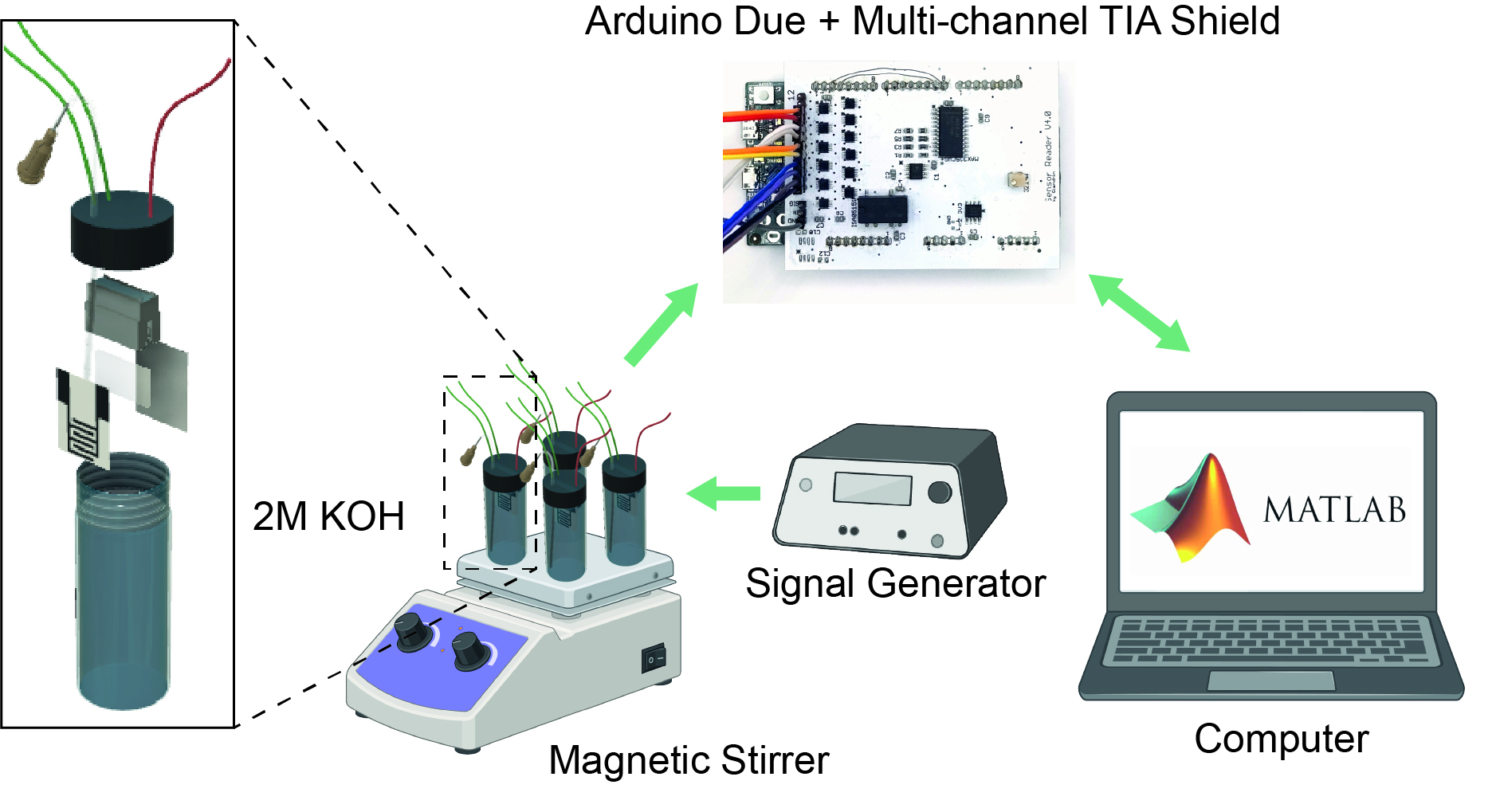

**Figure S4**. Setup for the sensor characterization experiment. Two chemPEGS were place on each glass vial and they were connected by an electrical feedthrough. The sensor signal is amplified and processed by the custom designed multi-channel transimpedance amplifier shield. The shield was connected to the Arduino Due for read-out and plotted on a computer through MATLAB.

**Calculation of rate of change in conductance of chemPEGS**

The rate of conductance change for the chemPEGS sensor was determined through a linear regression analysis performed on a specific segment of the data. To ensure consistency across experiments, the raw conductance data was first normalized. A baseline conductance (G_0_) was established by measuring the stable conductance of chemPEGS under fully humidified conditions where the liquid-vapor system was in equilibrium. All subsequent conductance measurements were then normalized relative to this G_0_ baseline.

A 3-minute analysis window was manually selected from a plot of the normalized conductance versus time. This window begins 15 seconds after the introduction of potassium hydroxide (KOH) to the sample solution. This specific interval was chosen because it corresponds to the period where the conductance changes at a constant rate. This change is driven by the neutralization of sulfuric acid on the sensor surface upon exposure to ammonia gas, which is liberated from the enzymatic breakdown of free asparagine. The slope derived from the linear regression of this 3-minute data segment represents the final rate of conductance change correlated to the free asparagine level in the sample solution.

**Linear Regression Analysis.** A linear regression model was applied to the normalized conductance data within selected time interval window. The analysis uses the standard linear equation ^1^:

$$\boldsymbol{y=a+bx}$$

Where,

**y** represents the normalized conductance of the chemPEGS.

**x** represents time.

**b** is the slope of the line, which directly quantifies the rate of change in conductance.

**a** is the y-intercept

The rate of change in conductance, **b** is calculated using the method of least squares.

Mathematically, **b** is calculated with the following formula ^2^:

$$\boldsymbol{b}=\frac{\sum_{i=1}^{n} x_{i}y_{i}-\frac{\left( \sum_{i=1}^{n} y_{i} \right)\left( \sum_{i=1}^{n} x_{i} \right)}{n}}{\sum_{i=1}^{n} x_{i}^{2}-\frac{\left( \sum_{i=1}^{n} x_{i} \right)^{2}}{n}}$$

Where:

- xᵢ and yᵢ are the individual time and conductance data points.
- x̄ and ȳ are the mean of the time and conductance values respectively

This calculation is performed automatically by regression functions of MATLAB to determine the precise rate of change in conductance of chemPEGS.

| Component | Cost (USD) |
| --- | --- |
| **Reusable Hardware (One-Time Cost)** | |
| 3D Printed Mini Magnetic Stirrer (Electronics + Hardware components) | $ 7.50 |
| Glass Vial + Lid | $ 2.52 |
| NFC PCB | $ 0.04 |
| SIC4341 IC Chip | $ 0.10 |
| Capacitors (x2) | $ 0.02 |
| **Total** | **$ 10.18** |
| **Consumable Components (Per-Test Cost)** | |
| L-Asparaginase | $ 1.26 |
| Potassium Hydroxide (KOH) | $ 0.11 |
| Sulfuric Acid (H_2_SO_4_) | <$ 0.01 |
| Paper-based Electrical Gas Sensor | <$ 0.01 |
| **Total** | **$ 1.39** |

**Figure S5**. The cost breakdown of the NFC-based FAsn sensing system in USD
